# Anterior cingulate cortex projections to the amygdala in primates: topographic and layer-specific organization underlying emotion and mood regulation

**DOI:** 10.64898/2026.02.09.704764

**Authors:** Kei Kimura, Rintaro Yoshino, Yuki Soga, Andi Zheng, Satoshi Nonomura, Gaoge Yan, Soshi Tanabe, Shinya Nakamura, Shinya Ohara, Ken-Ichi Inoue, Masahiko Takada, Ken-Ichiro Tsutsui

**Author notes:** Co-Corresponding authors: Masahiko Takada, Ken-Ichiro Tsutsui.

## Abstract

Emotion and mood regulation critically depends on interactions between the anterior cingulate cortex (ACC) and the amygdala. However, the detailed architecture of ACC projections to their major targets, the basal (BA) and accessory (AcBA) basal nuclei of the amygdala, remains unclear. To address this issue, a combined retrograde and anterograde tracing with viral vectors were performed in macaques to map the projection patterns from pregenual (pgACC), subgenual (sgACC), and dorsal (dACC) subareas. Data revealed that ACC neurons projecting to the BA arose predominantly from the superficial layers (II/III) of all subareas and the deep layers (V/VI) of the sgACC, whereas ACC neurons projecting to the AcBA originated mainly in the deep layers of the sgACC and dACC. The present study defines the topographic and layer-specific organization of ACC–amygdala connectivity in primates and subserves to provide an anatomical basis for future causal and translational approaches, such as targeted interventions against ACC-related mood disorders.

**Teaser:** Primate anterior cingulate cortex has topographic and layer-specific projections to amygdala that are involved in emotion and mood regulation.

## Introduction

Emotion and mood regulation is crucial for adaptive behavior, and its disruption causes serious mental illness, such as depression and anxiety disorders (for reviews, see 1–3). The regulatory mechanism underlying emotion and mood fundamentally relies on interactions between cortical and subcortical limbic structures. As widely recognized, the amygdala is largely involved in encoding negative emotion (4–7). While the amygdala is known to have neuronal connections with various cortical areas, the projections from the anterior cingulate cortex (ACC) to the amygdala are particularly important for the regulation of emotion and mood (6, 8, 9). To know about how emotion and mood are regulated, it is thus essential to elucidate the architecture of the neural pathway linking these two structures.

The ACC has long been considered a critical ‘hub’ for the information flow around the limbic system, including the amygdala (9–12). In humans and nonhuman primates (NHPs), the ACC is generally divided into structurally and functionally heterogeneous subareas, such as the pregenual, subgenual, and dorsal ACC (pgACC, sgACC, and dACC), which correspond respectively to Brodmann areas 32/24b, 25/14c, and 24a/b/c (13–15). Meta-analyses of numerous human neuroimaging studies indicate that these subareas of the ACC are involved in various aspects of emotion and mood regulation (13, 16). The importance of the ACC in the regulation of mood has also been highlighted in NHPs (17–18). Using repetitive transcranial magnetic stimulation, we have recently found that inhibitory interventions mainly targeting the pgACC and sgACC induce a typical depressive state in macaques (18).

There is a consensus that the ACC projections in NHPs are mainly directed toward the basal (BA) and accessory basal (AcBA) nuclei of the amygdala (6, 12, 19, 20). However, the origins of these projections within the ACC remain controversial. An early study (21) demonstrated that the ACC–amygdala projections arise from the sgACC (area 25) and dACC (dorsal area 24), whereas a more recent work (12) reported the existence of massive projections not only from the sgACC and dACC, but also from the pgACC (areas 32/24b). Therefore, a precise examination is needed for understanding the architecture of the ACC–amygdala pathway.

In the present study, we utilized retrograde/anterograde tracing approaches with adeno-associated virus (AAV) vectors to investigate the detailed neural connectivity of the ACC to the amygdala in macaques. For retrograde labeling from the amygdala, we injected a retrogradely transported AAV variant (so-called AAV2retro) into the BA and AcBA (22). To confirm the retrograde labeling data, we further conducted multi-color anterograde tract-tracing from the ACC by using a mosaic AAV vector (so-called AAV2.1; 23) that achieves high levels of transgene expression and neuron specificity. In a single animal, separate injections of AAV2.1 vectors expressing three different fluorescent proteins were made into the dACC, pgACC, and sgACC, respectively. These approaches indeed allowed us to define the topographic and layer-specific organization of distinct ACC projections to the BA and AcBA.

## Results

### Sites of AAV2retro injections in the AcBA and BA

To examine the distribution of neurons in the medial frontal cortex (MFC) giving rise to the ACC–amygdala pathway, we injected AAV2retro into the AcBA or BA in four monkeys (Monkeys A-D; Fig. 1). In each case, the extent of the injection site was reconstructed in coronal sections at five anteroposterior levels selected to cover the whole region of the amygdala (Fig. 1a). In Monkeys A and B, AAV2retro injections were made into the AcBA (Fig. 1b). The injection site in each case was almost limited to the AcBA as involving its magnocellular (AcBAmc) and parvicellular (AcBApc) divisions, with minimal diffusion into the anterior hypothalamic area (Fig. 1a). In Monkeys C and D, on the other hand, AAV2retro was injected into the BA (Fig. 1b). The injection site in each case largely involved the magnocellular (BAmc) and intermediate (BAi) divisions, but not the parvicellular division of the BA, though slightly encroached on the lateral nucleus of the amygdala (Fig. 1a).

**Fig. 1.**
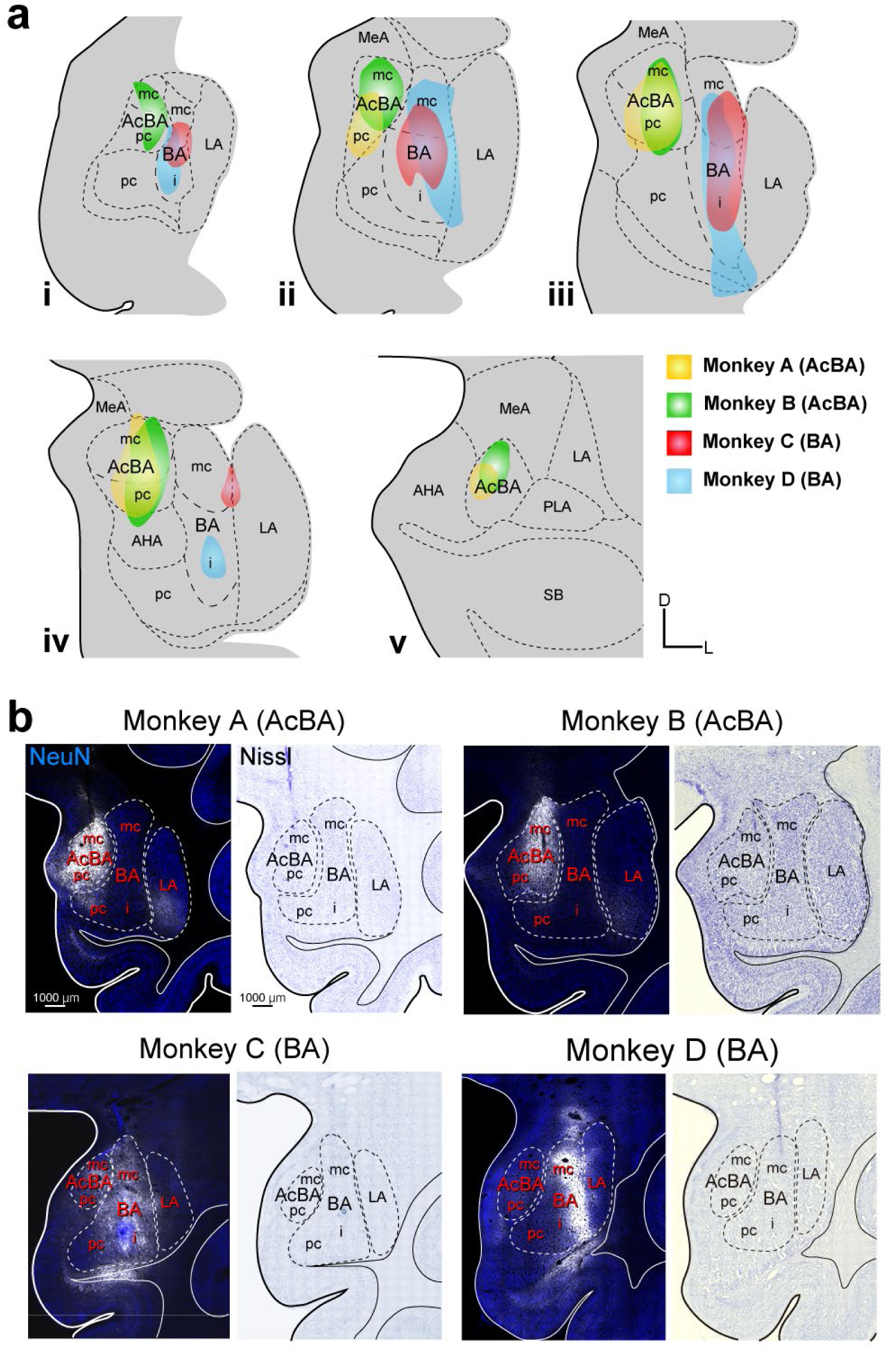
Sites of AAV2retro injections in the amygdala. a: Extent of the injection sites in the AcBA (Monkeys A and B) and BA (Monkeys C and D). Five representative coronal planes are arranged anteroposteriorly from (i) to (v) and the four injection cases are overlaid thereon. Regions in yellow and green show the spread of the injected vector within the AcBA in Monkeys A and B, respectively, and those in red and blue depict the spread of the injected vector within the BA in Monkeys C and D, respectively. b: Fluorescent images of the injection sites (left) and Nissl-stained cytoarchitecture (right) in Monkeys A-D. mKO2 native-fluorescence is shown in white, and NeuN immunofluorescence is shown in blue. AcBA, accessory basal nucleus of the amygdala; AHA, anterior hypothalamic area; BA, basal nucleus of the amygdala; D, dorsal; i, intermediate division; L, lateral; LA, lateral nucleus of the amygdala; mc, magnocellular division; MeA, medial amygdaloid area; pc, parvocellular division; PLA, paralaminar nucleus of the amygdala; SB, subiculum.

### Cells of origin of ACC projections to the AcBA and BA

To compare the distribution of cells of origin of ACC projections to the AcBA and BA, we plotted retrogradely labeled neurons in the MFC on coronal sections and prepared unfolded maps (see Materials and Methods). Consistent with previous findings (21), retrograde neuron labeling prominently occurred not only in the MFC, but also in the orbitofrontal cortex in all cases (Figs. S1 and S2). Within the MFC, more intense labeling was seen more posteriorly, especially in areas 25 and 14c corresponding to the sgACC in the four cases (Figs. 2, S1, and S2). However, the distribution patterns of labeled neurons in other MFC areas varied depending on the site of AAV2retro injection, either the AcBA or the BA.

**Fig. 2.**
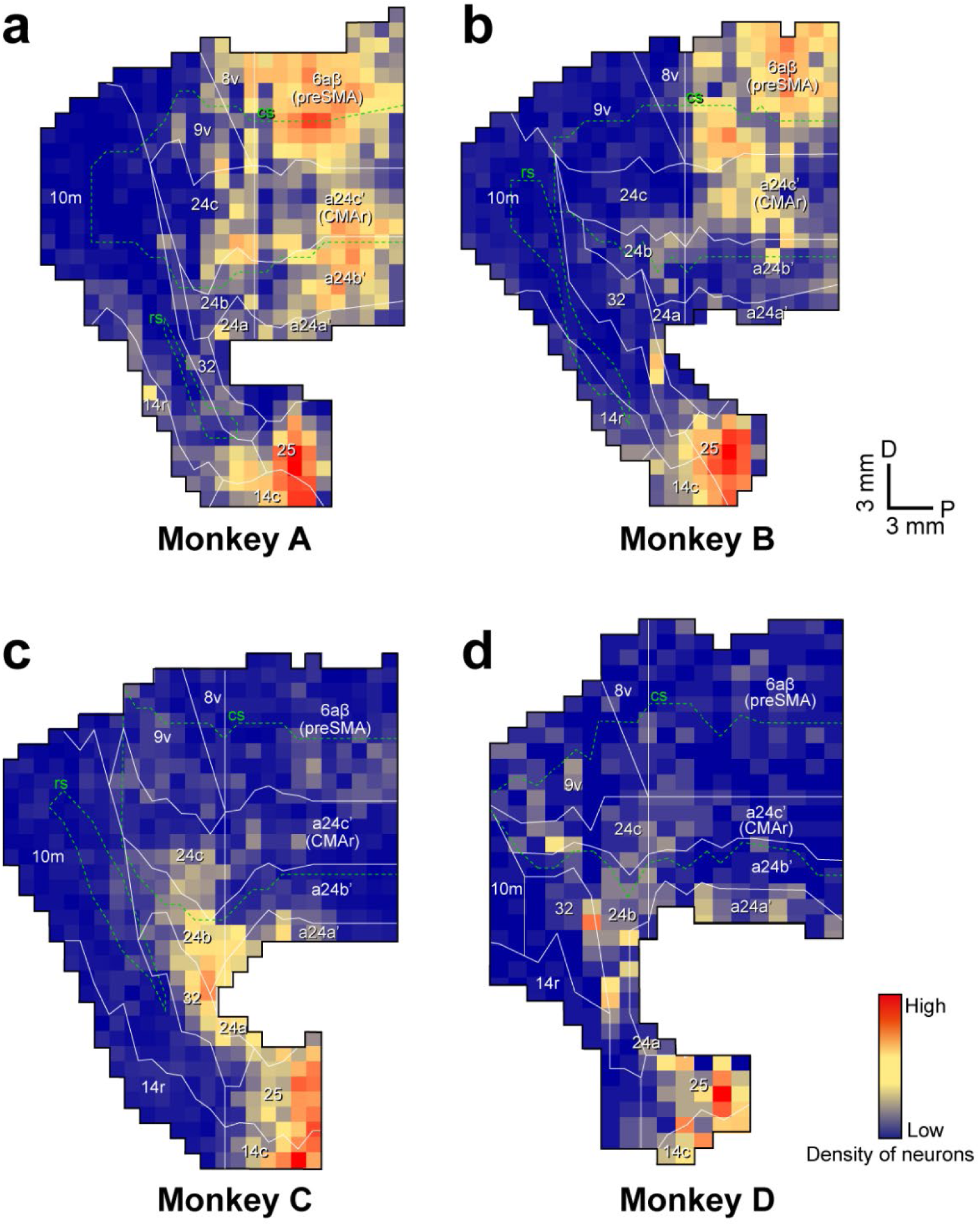
Unfolded maps of the MFC showing the distribution patterns of retrograde labeling from the AcBA in Monkeys A (a) and B (b) and from the BA in Monkeys C (c) and D (d) For each case, cell counts were performed at 1-mm intervals and normalized to the mean of all cell number within each section. In each 1-mm bin, the density of labeled cortical neurons is expressed as a colormap (red, high; yellow, intermediate; blue, low). CMAr, rostral cingulate motor area; cs, cingulate sulcus; D, dorsal; P, posterior; preSMA, pre-supplementary motor area; rs, rhinal sulcus.

In Monkeys A and B where the AAV2retro injections were made into the AcBA, dense clusters of labeled neurons were located in the sgACC (i.e., areas 25 and 14c), as described above (Figs. 2 and S1). Moreover, large numbers of labeled neurons were found in the posterior and dorsal part of the MFC, such as posterior areas 24b/c, the rostral cingulate motor area, and the pre-supplementary motor area (Figs. 2 and S1). A small number of neurons were also labeled in area 24a. Similar patterns of MFC labeling were observed in the two cases, although neurons in the posterior and dorsal MFC were more numerously labeled in Monkey A (Fig. 2).

In Monkeys C and D where the AAV2retro injections were performed into the BA, a large sector over the pgACC and dACC, including posterior area 32 and anterior areas 24a/b/c, contained clusters of labeled neurons (Figs. 2 and S2). Dense neuron labeling was seen in the sgACC (i.e., areas 25 and 14c) like the AcBA-injection cases, while unlike the AcBA-injection cases, only a few neurons were retrogradely labeled in the posterior and dorsal MFC (Figs. 2 and S2). Substantially the same patterns of MFC labeling were obtained in the two cases, though the labeled neurons in the pgACC and sgACC were more frequently found in Monkey C where the injection volume of AAV2retro was twice as much as in Monkey D (Figs. 2 and 4).

### Differential layer distributions of ACC neurons projecting to the AcBA vs. BA

Furthermore, we analyzed the layer distribution of ACC neurons projecting to the AcBA and BA. Unfolded maps prepared based on the ratio of superficial vs. deep layers neurons (see Materials and Methods) revealed that ACC neurons retrogradely labeled from the AcBA were primarily distributed within the deep layers (i.e., layers V and VI) throughout the MFC (Figs. 3a,b, 4a,b, and 5a), which is consistent with previous findings (21). By contrast, many neurons labeled from the BA were located within the superficial layers (i.e., layers II and III). Notably, the vast majority of labeled neurons in the pgACC (i.e., posterior area 32 and anterior area 24b) were observed in the superficial layers (Figs. 3c,d, 4c,d, and 5b). In addition, at least part of the sgACC (i.e., areas 25 and 14c) and dACC (i.e., anterior areas 24a/b/c) often contained labeled neurons in the superficial layers (Figs. 3c,d, 4c,d, and 5b).

**Fig. 3.**
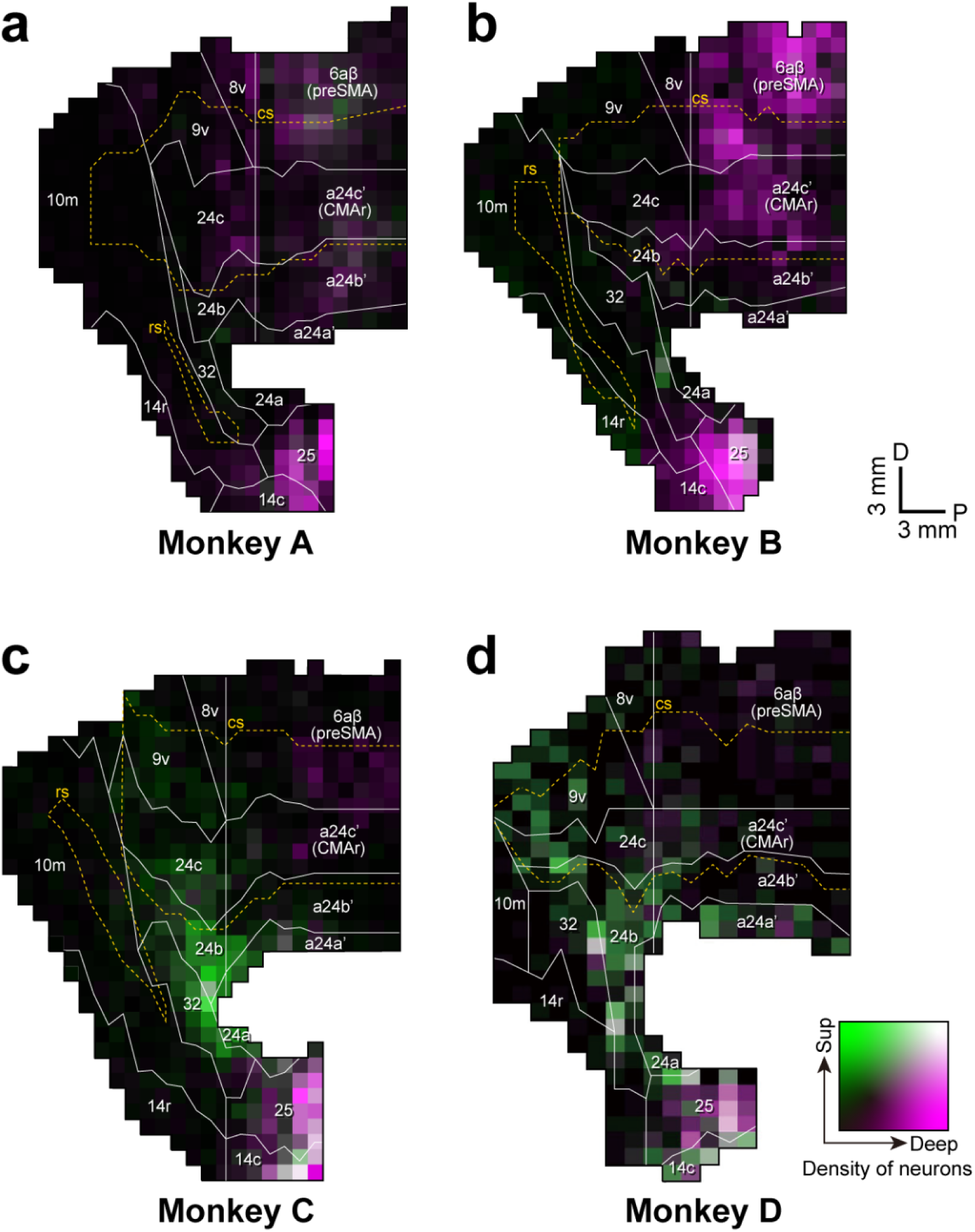
Unfolded maps of the MFC showing the layer distribution patterns of retrograde labeling from the AcBA in Monkeys A (a) and B (b) and from the BA in Monkeys C (c) and D (d) For each case, cell counts were performed at 1-mm intervals and normalized to the mean cell count. Each 1-mm bin is indicated with a bivariate colormap encoding superficial (layers II/III) versus deep (layers V/VI) layer densities: Bins enriched in superficial-layer labeling are denoted in green; those enriched in deep-layer labeling in magenta; those with high densities in both layers in white; and those with low densities in both layers in black. Deep, deep layers; Sup, superficial layers. Other abbreviations are as in Figure 4.

**Fig. 4.**
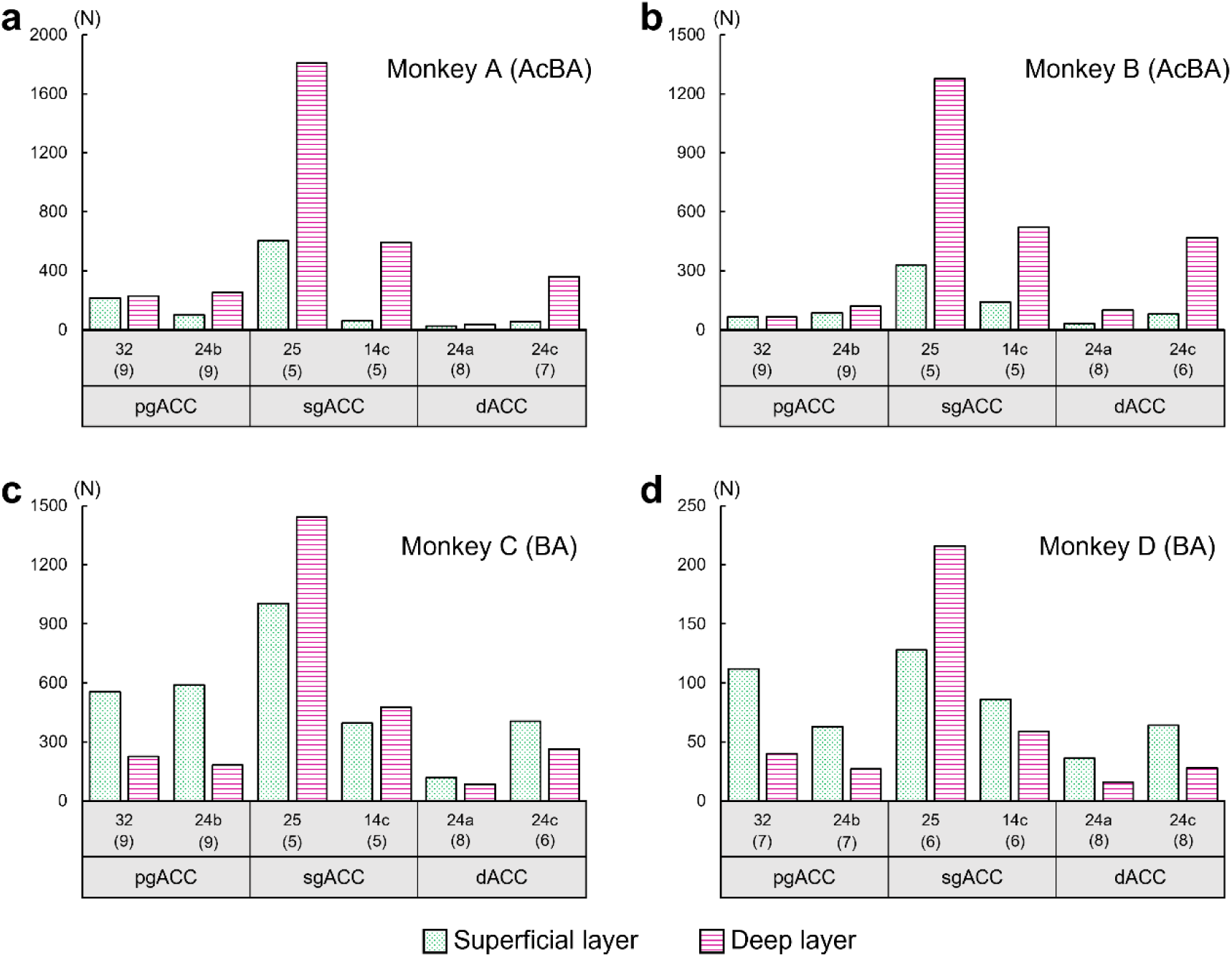
Quantification of ACC neurons projecting from the superficial vs. deep layers to the AcBA and BA (a,b) Number of labeled neurons projecting from the pgACC, sgACC, and dACC to the AcBA in Monkeys A (a) and B (b). (c, d) Number of labeled neurons projecting from the pgACC, sgACC, and dACC to the BA in Monkeys C (c) and D (d). Green dots represent neurons projecting from the superficial layers, while magenta lines represent neurons projecting from the deep layers. The number of coronal sections used for cell counts is indicated in parentheses under each cortical area number.

**Fig. 5.**
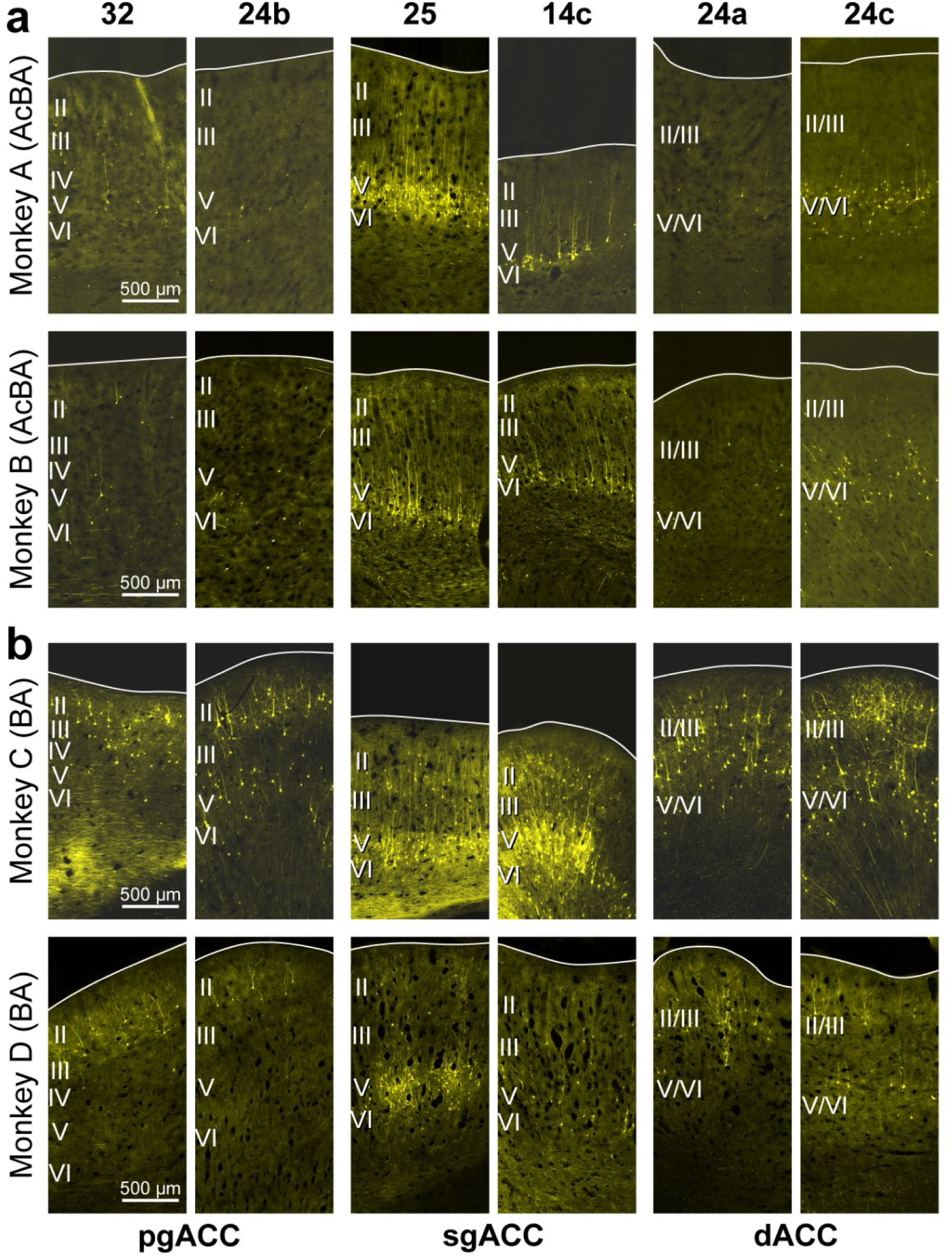
Fluorescent images showing the layer distribution patterns of retrograde labeling from the AcBA in Monkeys A and B (a) and from the BA in Monkeys C and D. **(b)** Fluorescent photomicrographs taken from the pgACC (areas 32 and 24b), sgACC (areas 25 and 14c), and dACC (areas 24a and 24c). The layer assignment was cytoarchitectonically determined based on adjacent Nissl-stained sections. Note that the projections from the pgACC and dACC to the BA arise predominantly from the superficial layers (II/III), whereas the other projections arise mainly from the deep layers (V/VI).

### Topographic distributions of ACC-derived terminals in the AcBA and BA

To confirm the data obtained from the retrograde labeling experiments, multi-color anterograde tract-tracing was carried out by using three AAV2.1 vectors expressing three different fluorescent proteins. In Monkey E, these vectors were injected separately into the pgACC, sgACC, and dACC (Fig. 6a,b). The pgACC injection was placed in area 32 and, also, rostral area 24b, and produced dense terminal labeling in the BA, especially in the BAi (Figs. 6 and 7). Some labeled terminals were further detected in the BAmc and AcBAmc. The sgACC injection was placed in areas 25 and 14c, with slight diffusion into ventral area 32 and area 24a. After the sgACC injection, anterogradely labeled terminals were distributed widely in both the BA and the AcBA, with particular dominance in the BAi and AcBAmc (Figs. 6 and 7). Following the dACC injection covering anterior areas 24a/b/c, terminal labeling was seen sparsely but extensively over the BA (including both the BAmc and the BAi) and AcBA (including both the AcBAmc and the AcBApc) (Figs. 6 and 7).

**Fig. 6.**
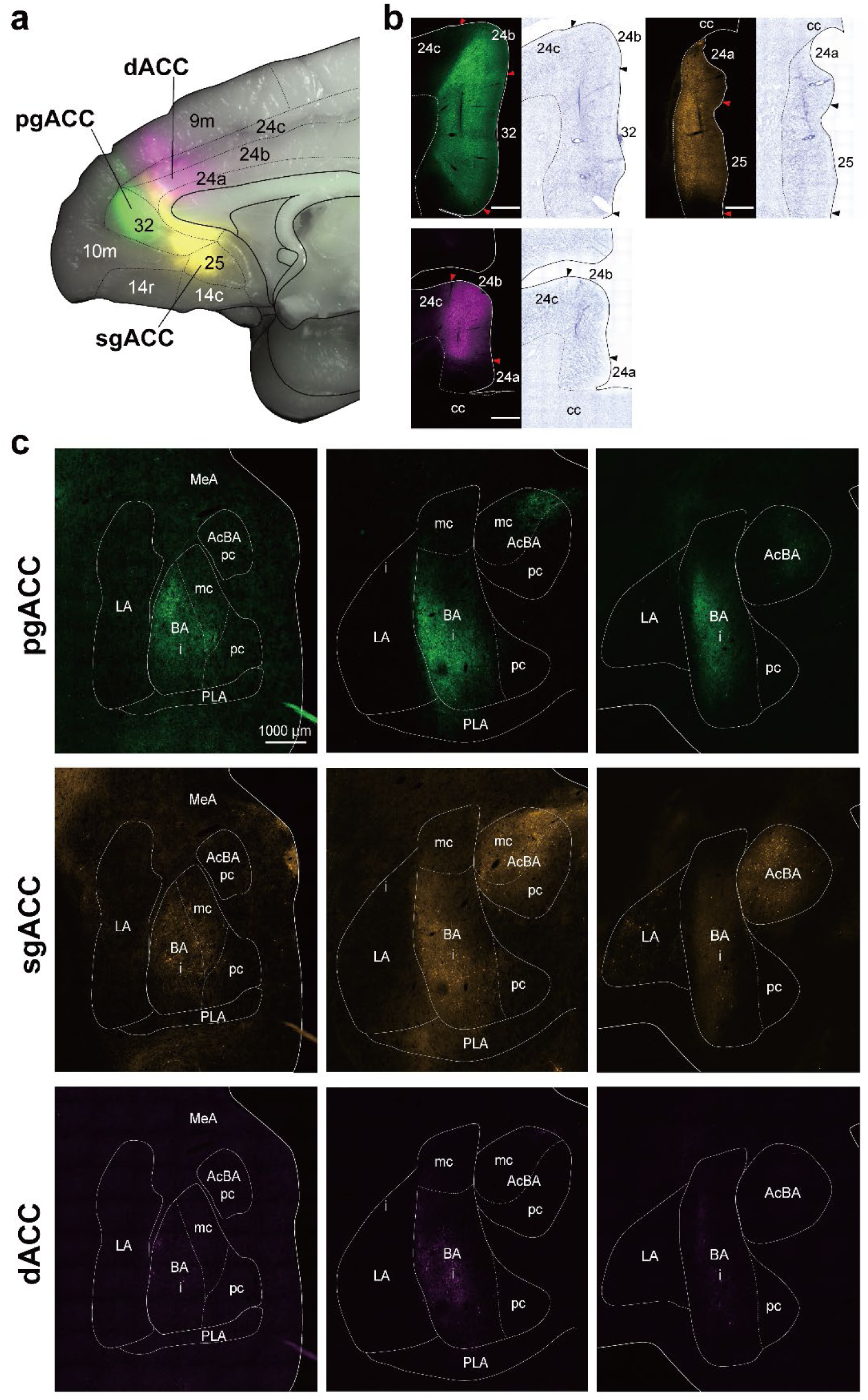
Anterograde labeling in the amygdala after AAV2.1 injections into the pgACC, sgACC, and dACC in Monkey E (a) Low-power image of the injection sites of three vectors (AAV2.1-CaMKII-mClover, AAV2.1-CaMKII-mKO2, and AAV2.1-CaMKII-mRaspberry) in the pgACC (green), sgACC (orange), and dACC (magenta). (b) Representative coronal planes through the injection sites in the pgACC (green), sgACC (orange), and dACC (magenta) in conjunction with the Nissl-stained cytoarchitecture. (c) Arranged anteroposteriorly are three representative coronal planes through the amygdala following the pgACC injection (top), sgACC injection (middle), and dACC injection (bottom). Anterogradely labeled terminals from the pgACC, sgACC, and dACC are shown in green, orange, and magenta, respectively. cc, corpus callosum. Other abbreviations are as in Figure 1.

**Fig. 7.**
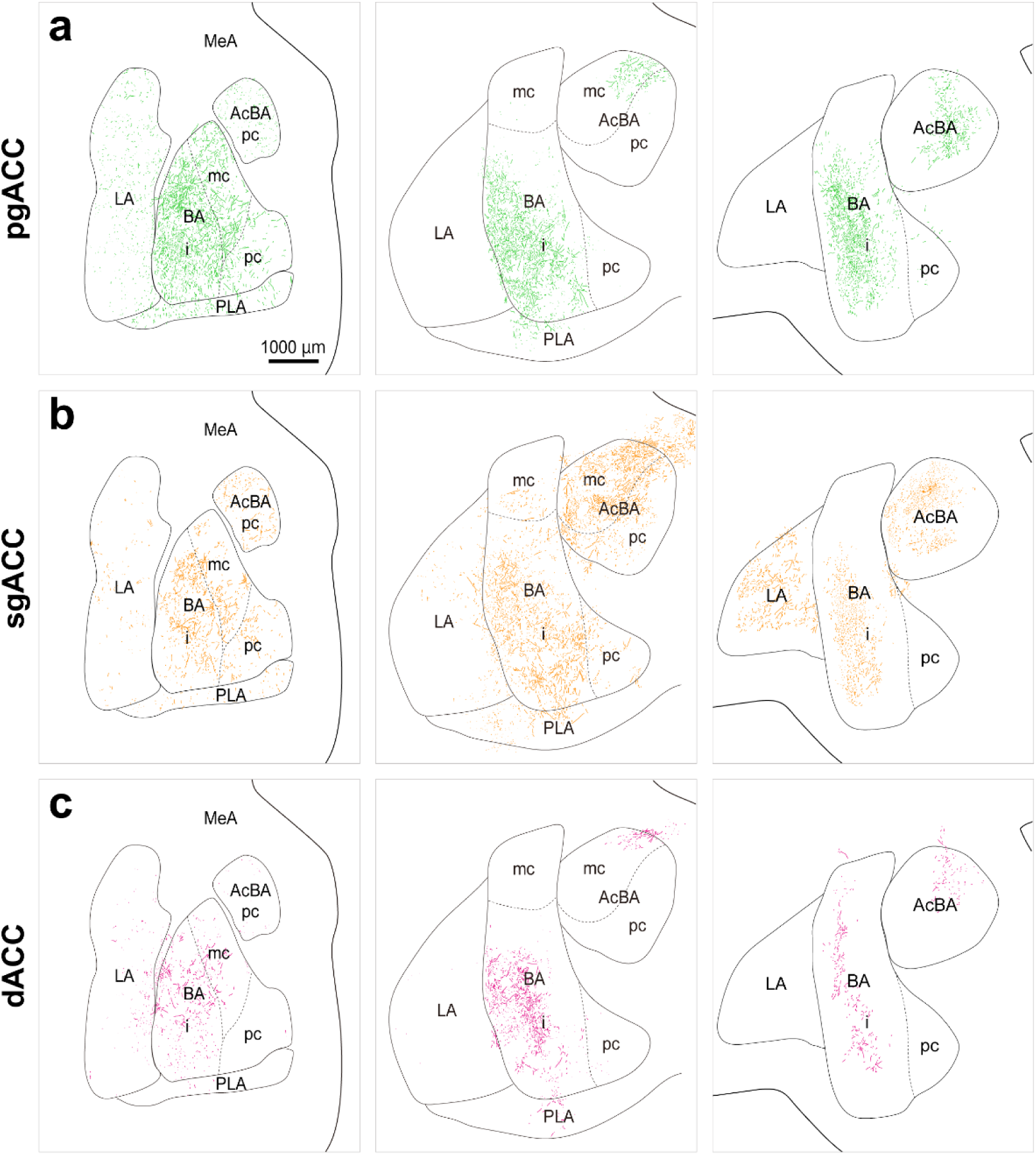
Distribution patterns of anterograde labeling in the amygdala after AAV2.1 injections into the pgACC (a), sgACC (b), and dACC (c) In each case, three representative coronal planes through the amygdala are arranged anteroposteriorly. Anterogradely labeled terminals from the pgACC, sgACC, and dACC are plotted in green, orange, and magenta, respectively. All abbreviations are as in Figure 1.

## Discussion

In the present study, AAV vector systems permitting sensitive retrograde and anterograde neural tracing were employed to investigate precisely the architecture of ACC–amygdala projections in macaques. We have identified that the three distinct areas of the ACC, the pgACC, sgACC, and dACC, exhibit different projection patterns to the BA and AcBA (Fig. 8). As indicated by the results on our retrograde viral tracing, although the pgACC, sgACC, and dACC each possess widespread projections to the BA, projections to the AcBA arise mainly from the sgACC, with only sparse input originating in the pgACC. This projection pattern to the AcBA is consistent with the previous report (21), and our findings further support the existence of a novel projection from the pgACC to the BA, as also described in the prior work (12). Because the BA has a limited extent along the anteroposterior and mediolateral axes, achieving a highly localized tracer injection to the BA is technically challenging. Consequently, many previous studies that targeted the BA with retrograde tracers likely failed to confine their injections to the BA and included some degree of leakage into the AcBA, potentially leading to an underestimation of pgACC-derived projections relative to those from the sgACC and dACC.

**Fig. 8.**
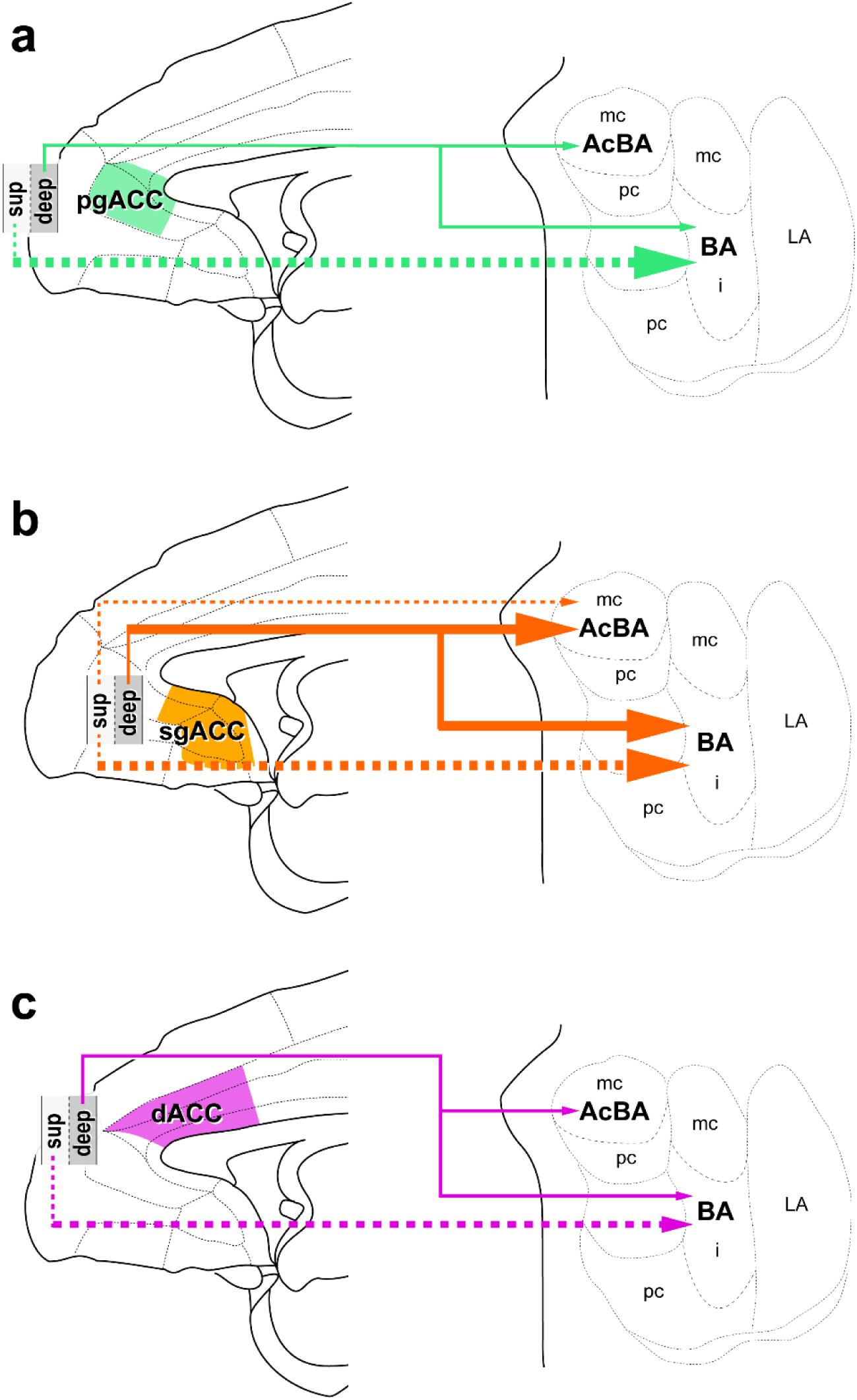
Schematic diagram showing the topographic and layer-specific projections from the ACC to the amygdala Projections from the pgACC (a; green), sgACC (b; orange), and dACC (c; magenta) to the BA and AcBA. Superficial and deep layers projections are depicted by dotted and solid lines, respectively. Arrow width represents the intensity of projections. All abbreviations are as in Figure 1.

These NHP data provide an important clue to considering the functional parcellation within the MFC and encourage reconsideration of a prevailing rodent model. In rodents, it has been shown that MFC subregions exhibit the target- and layer-dependent differences in their projections to the amygdala and other limbic and autonomic centers (24, 25). More recent works have also demonstrated that the prelimbic cortex (PL) projects primarily to the basolateral amygdala (26), whereas the infralimbic cortex (IL) projects mainly to the basomedial amygdala (26, 27). Based on such an anatomical framework, several early studies in rodents proposed a relatively simple functional dissociation within the MFC, in which the PL facilitates fear expression whereas the IL promotes the consolidation and recall of fear extinction (28–32).

However, accumulated evidence to date suggests that this dichotomy may be simplistic. Notably, neuronal activity in both the PL and the IL increases during the recall of fear extinction, i.e., the memory of safety (33, 34), indicating that these regions may contribute jointly, and in a task-dependent manner, to extinction-related processes rather than supporting strictly opposing functions. These complexities become even more apparent when considering findings from human studies. Structural and functional imaging studies in individuals with major depressive disorder consistently report gray matter reduction and hyperactivity in the sgACC (14, 35, 36). fMRI works have further demonstrated dissociable contributions of the pgACC and sgACC to affective processing (16, 37): the pgACC is implicated in fear extinction and stress resilience (38–40), whereas the sgACC is associated with negative affective processing and depressive symptomatology (13, 37, 41, 42). These observations in humans do not clearly align with the functional distinctions proposed in early rodent studies, resulting in a fragmented and often inconsistent understanding of how MFC–amygdala circuits contribute to affective regulation across species. However, human neuroimaging studies alone cannot establish the circuit-level mechanisms underlying these affective functions. The lack of cross-species correspondence highlights the need for studies in NHPs whose MFC–amygdala organization more closely resembles that of humans. Recent causal manipulations in NHPs have begun to address this gap. We have recently reported that low-frequency repetitive transcranial magnetic stimulation (rTMS) targeting the monkey MFC, which was centered on the pgACC and sgACC, induces depressive-like phenotypes (18). Complementing this approach, deep brain stimulation near the subcallosal ACC has likewise produced large-scale network alterations in macaques (43), reinforcing the value of primate models for probing the causal mechanisms of affective dysfunction. Nonhuman primate models of mood disorders often recapitulate key clinical features observed in humans, underscoring their value for bridging mechanistic gaps that remain unresolved in rodent studies.

Consequently, detailed investigations of limbic circuit architecture in monkeys are increasingly needed to advance our understanding of the neural mechanisms underlying psychiatric disorders, which remain largely enigmatic.

Importantly, our cortical laminar distribution analysis has revealed a marked contrast between the ACC projections to the BA and the AcBA: whereas the inputs to the AcBA arise almost exclusively from the deep layers, those to the BA exhibit the substantial involvement of the superficial layers (see Figs. 3–5, and 8). Notably, among the ACC subareas, only the pgACC showed a predominance of the superficial-layer projections over the deep-layer projections in its input to the BA. Cortical outputs to subcortical structures typically arise from neurons in the deep layers, whereas superficial-layer neurons primarily contribute to corticocortical pathways (44).

Thus, superficial-layer projections to subcortical targets are considered highly exceptional. Interestingly, however, similar superficial-layer projections from the MFC to the amygdala have also been reported in rodents (25, 45, 46), suggesting that this laminar feature may represent a conserved but underappreciated aspect of the MFC–amygdala connectivity across species. The ACC is further notable for exhibiting marked developmental transitions in cellular and circuit organization. In rodents, microglial morphology and activity in the MFC change dynamically throughout the developmental stages (47, 48). Consistent with this, our previous work has shown coordinated activation of astrocytes, microglia, and oligodendrocyte-lineage cells in the macaque ACC during late infancy and puberty (49). Moreover, it has recently been demonstrated that some developmental changes occur in the functional impact of MFC projections to the basolateral amygdala in rodents (50). In addition, transcriptomic analyses in primates have revealed substantial shifts of gene expression in the ACC during adolescence (51, 52). Together, these findings raise the possibility that the superficial-layer ACC projections to the BA emerge as part of broader developmental reorganization of the limbic-related circuitry. Although the precise function of the pgACC superficial-layer projections to the amygdala remains to be determined, the ACC receives strong cognitive inputs from the prefrontal cortical areas (53–55). One speculative possibility is that superficial-layer pgACC neurons contribute to the maturation of cognitive control over emotion (including resilience) by providing a developmentally refined route through which higher-order cognitive signals can modulate information processing in the amygdala. A recent theoretical work also highlights the contribution of the ACC to predictive coding integrating cognitive and emotional information, suggesting that the ACC–amygdala pathway may support predictions that guide affective responses (56). In this context, the superficial-layer-derived pathway may be responsible for more rapid or efficient transfer of cognitive information from the prefrontal cortex to the amygdala, thereby subserving the effective integration of cognitive and affective processes in the mature brain.

To further advance our understanding of the ACC–amygdala circuitry, pathway-selective causal manipulations should be called for. In particular, separately manipulating the activity of amygdala-projecting neurons arising from the superficial vs. deep layers of the ACC subareas will help to dissociate their specific functions. Prior studies on the monkey ACC indicate its extensive roles not only in affective responses, but also in decision-making and social cognition (57–63).

Therefore, layer-selective manipulations of the ACC–amygdala pathways in macaques by means of optogenetic/chemogenetic techniques may enable us to define individual pathways critical for these cognitive processes (64). Considering the developmental changes highlighted above, future works should take into account the possibility that these pathways undergo substantial reorganization during postnatal maturation. Developmentally informed circuit analyses, including comparisons of the laminar origins, synaptic targets, and functional roles of the ACC–amygdala projections throughout the juvenile, adolescent, and adult stages, will be essential for determining whether the superficial- and deep-layer pathways acquire differential functions as the ACC matures. In addition to layer- and pathway-level distinctions, the ACC itself exhibits pronounced neurochemical heterogeneity across its subareas. Multi-receptor autoradiography in the human cingulate cortex has demonstrated that the ACC subareas such as the pgACC and sgACC differ markedly in their receptor fingerprints, including AMPA, kainate, GABAB, and 5-HT1A receptor densities (65). These findings suggest that the ACC subareas are configured with distinct neurochemical architectures that may interact with their layer-specific projection patterns to shape affective and cognitive functions. Incorporating such receptor-level information into future circuit analyses will be important for understanding how developmental or pathological perturbations selectively impact the ACC–amygdala pathways. Parsing the ACC circuitry at this finer spatial and temporal granularity may ultimately provide critical insights into the mechanisms by which developmental perturbations contribute to mood disorders and may help to guide the design of targeted interventions against the ACC-related dysfunctions.

## Materials and Methods

### Animals and Experimental Design

Five adult macaques were used for this study (Monkeys A-E in Table 1): four rhesus monkeys (*Macaca mulatta*), 5.7-19.5 years old, two males and two females, 5.0-8.1 kg; one Japanese monkey (*Macaca fuscata*), 7.2 years old, male, 9.8 kg. They were housed in individual cages (approximately W750×D900×H900 mm) in a room with controlled temperature (23-26℃) and light (12-hr on-off cycle) conditions. Every effort was made to minimize animal suffering.

**Table 1.** Summary of Experiments.

| Animal ID | Sex | B.W.<br>(kg) | Age<br>(yo) | Target | Serotype | Promoter | Gene | Injection<br>titer<br>(gc/ml) | Injection<br>volume<br>(µl/target) |
| --- | --- | --- | --- | --- | --- | --- | --- | --- | --- |
| Monkey A | M | 8.1 | 6.7 | AB | AAV2retro | hSyn | mKO2 | 2.0×10<sup>13</sup> | 4.8 |
| Monkey B | F | 6.1 | 5.7 | AB | AAV2retro | hSyn | mKO2 | 2.0×10<sup>13</sup> | 4.5 |
| Monkey C | M | 6.9 | 19.5 | B | AAV2retro | hSyn | mKO2 | 2.0×10<sup>13</sup> | 4.8 |
| Monkey D | M | 9.8 | 7.2 | B | AAV2retro | hSyn | mKO2 | 2.0×10<sup>13</sup> | 2.4 |
| Monkey E | F | 5.0 | 9.6 | pgACC | AAV2.1 | CaMKII | mClover | 2.0×10<sup>13</sup> | 2.0 |
|  |  |  |  | sgACC | AAV2.1 | CaMKII | mKO2 | 2.0×10<sup>13</sup> | 2.0 |
|  |  |  |  | dACC | AAV2.1 | CaMKII | mRaspberry | 2.0×10<sup>13</sup> | 2.0 |

B.W., body weight; hSyn, human synapsin promoter.

In Monkeys A and B, AAV2retro was injected unilaterally into the AcBA, with two tracks delivered in each monkey (Table 1). In Monkeys C and D, AAV2retro injections were also made unilaterally into the BA, with one track delivered in Monkey C and two tracks delivered in Monkey D (Table 1). In Monkey E, on the other hand, three AAV2.1 vectors expressing distinct fluorescent proteins were injected separately into the pgACC, sgACC, and dACC of the same hemisphere (Table 1). According to preceding studies (13–15), we cytoarchitectonically defined the pgACC as Brodmann areas 32 and 24b, the sgACC as areas 25 and 14c, and the dACC as areas 24a, 24b, and 24c.

The entire research project was approved by the Tohoku University Animal Care and Use Committee (Tohoku University) and the Animal Welfare and Animal Care Committee of the Center for the Evolutionary Origins of Human Behavior (Kyoto University). All experiments were conducted in accordance with the Regulations for Animal Experiments and Related Activities at Tohoku University (Version 18, 2025), the Guidelines for Care and Use of Nonhuman Primates established by the Primate Research Institute, Kyoto University (2010), and the Guide for the Care and Use of Nonhuman Primates in Neuroscience Research (Japan Neuroscience Society).

### Surgery

The monkeys were initially sedated with ketamine hydrochloride (5 mg/kg, i.m.) and xylazine hydrochloride (0.5 mg/kg, i.m.), and then anesthetized with sodium pentobarbital (20 mg/kg, i.v.) or propofol (5 mg/kg/hr). An antibiotic (Ceftazidime; 25 mg/kg, i.v.) was administered at the initial anesthesia. During the surgery, the monkeys were kept hydrated with a lactated Ringer’s solution (i.v.) and, also, applied with the same antibiotic (Cefazolin; 250-500 mg/kg, i.m.) and an analgesic (Buprenorphine hydrochloride; 0.02 mg/kg, i.m.). After removal of a skull portion over the frontal lobe, stereotaxic injections of AAV vectors were made by the aid of a magnetic resonance imaging (MRI)-guided navigation system (Brainsight Primate; Rogue Research, Montreal, Canada). The AAV2retro expressing red fluorescent protein (RFP) (AAV2retro-hSyn-mKO2) was injected unilaterally into the AcBA or BA, while the AAV2.1 vectors expressing three different fluorescent proteins (AAV2.1-TetOff-CaMKII-pal-mClover/mKO2/mRaspberry) were injected, respectively, into the pgACC, sgACC, and dACC of the same hemisphere (Table 1). The AAV2.1 vector is a mosaic capsid vector that exhibits high neuronal specificity and robust transgene expression in the primate brain (23). In the present study, we incorporated a tetracycline transactivator (tTA) gene downstream of the mouse CaMKIIα promoter and TRE-tight promoter upstream of the fluorescent genes to enhance transgene expression, thereby facilitating the visualization of fine axon terminals (66–68). The details of production for AAV vector were mentioned elsewhere (23). All injections were performed through a 10-μl Hamilton Neuros Syringe (Hamilton, Reno, USA) at the rate of 0.1-0.2 μl/min. The injection volume of AAV2retro in Monkeys A-D was 2.0-2.5 μl/site (one site per track; one or two tracks), and that of AAV2.1 vectors in Monkey E was 0.5 μl/site (two sites per track; two tracks). The injection titer for each vector is provided in Table 1. After the injections, the monkeys were monitored until full recovery from the anesthesia. All experimental procedures were carried out in a special laboratory designated for *in vivo* animal infectious experiments (biosafety level 2 at Kyoto University and level 1 at Tohoku University).

### Tissue Preparation

Three to four weeks after the vector injections, the monkeys underwent perfusion-fixation. Each animal was anesthetized deeply with an overdose of sodium pentobarbital (50 mg/kg, i.v.), sodium secobarbital (50 mg/kg, i.v.), or sodium thiopental (100 mg/kg, i.v.), and perfused transcardially with 0.1 M phosphate-buffered saline (PBS; pH 7.4), followed by 10% formalin in 0.1 M phosphate buffer (pH 7.4). The removed brain was postfixed in the same fresh fixative overnight at 4°C and then saturated with 30% sucrose in 0.1 M PBS at 4°C. Coronal sections were cut serially at the 50-µm thickness on a freezing microtome and divided into ten series.

### Histological analyses

Every tenth section 500-µm apart was mounted onto glass slides and observed under a fluorescence microscope (AxioScan Z1; Carl Zeiss, Oberkochen, Germany). For cytoarchitectonic delineation of cortical areas/layers and amygdala nuclei, the adjacent series of sections were mounted onto gelatin-coated glass slides and Nissl-stained with 1% Cresyl violet. To confirm the distinction of injection sites, the fluorescence histochemistry for mClover, mKO2, mRaspberry, and NeuN was performed for Monkey E. First, Coronal sections were rinsed three times in 0.1 M PBS, and immersed in 1% skim milk for 1 hr. These sections were then incubated for two days at 4 °C with a cocktail of rabbit monoclonal antibody against mClover (1:2000; Invitrogen, Waltham, USA), mouse monoclonal antibody against mKO2 (1:750; Medical & Biological Laboratories, Nagoya, Japan), rat monoclonal antibody against mRaspberry (1:500; ChromoTek, Martinsried, Germany), and guinea pig polyclonal antibody against NeuN (1:1,000; Millipore, Billerica, USA) in 0.1 M PBS containing 2% normal donkey serum and 0.1% Triton X-100. Subsequently, the sections were incubated with a cocktail of Alexa 488-conjugated donkey anti-rabbit IgG antibody (1:200; Jackson ImmunoResearch, West Grove, USA), Alexa 555-conjugated donkey anti-mouse IgG antibody (1:200; Invitrogen), Alexa 647-conjugated donkey anti-rat IgG antibody (1:200; Jackson ImmunoResearch), and DyLight 405-conjugated donkey anti-guinea pig IgG antibody (1:200; Jackson ImmunoResearch) in the same fresh medium for 2 hr at room temperature. In addition, NeuN immunohistochemistry was performed for Monkeys A–D using the same procedure. All histological sections were mounted onto gelatin-coated glass slides. NeuN immunofluorescence images were not included in the figures, but were used solely to identify the regional boundaries during histological analysis.

In Monkeys A-D, the number of RFP-positive neurons was counted in the MFC. Based on these data, unfolded maps of the MFC were constructed from coronal sections 1-mm apart (Figs. 2 and 3). To quantify the density of neurons in individual areas of the MFC, we performed an unfolded map analysis, in which cortical sulci were virtually unfolded and the cortical sheet was flattened into a two-dimensional plane to visualize the distribution of labeled neurons (69).

Coronal-section images obtained at 1-mm intervals were imported into Adobe Illustrator (Adobe Inc., San Jose, USA), and regions of interest (ROIs) were manually delineated along the medial cortical surface. Each ROI was subdivided into bins extending from the dorsal to the ventral edge of the cortex. The bin boundaries were defined such that the length of the line separating layers V and VI was set to 1,000 µm, resulting in bins of trapezoidal or fan-like shapes. For the whole cell-count analysis without layer distinction, the number of labeled neurons within each bin was quantified from dorsal to ventral. For the cell-count analysis with layer distinction, labeled neurons were counted separately in superficial-layer bins (layers I–IV) and deep-layer bins (layers V and VI). Cell counts from each bin were entered into a matrix, with each column representing one coronal section arranged from anterior to posterior. The resulting matrix was then converted into a heatmap, in which color values were assigned based on normalization to the average number of labeled neurons across all bins. In addition, we assessed the layer-specific distribution patterns of labeled neurons. For this purpose, the superficial (layers II and III) and deep (layers V and VI) layers were separately subdivided into columnar bins (1,000-µm width), and the number of labeled neurons in each bin was counted. The bin-wise counts were then normalized to the total number of labeled neurons in the MFC, and the normalized values were plotted onto the unfolded maps to visualize the layer distribution patterns. To prepare Figure 2, the values for the superficial and deep layers were combined into single maps.

## Acknowledgments

We are grateful to M. Fujiwara, M. Nakano, E. Sumiya, E. Tanaka, W. Lu, X. Zhao and T. Moritani for their technical assistance.

## Author contributions

Conceptualization, M.T., and K-I.T.; formal analysis, K.K., R.Y., and Y.S.; funding acquisition, K.K., K-I.I., M.T., and K-I.T.; investigation, K.K., R.Y., Y.S., A.Z., S.No., G.Y., S.T., S.Na., S.O., K-I.I.; project administration, K-I.T; resources, K-I.I., M.T., and K-I.T.; supervision, M.T. and K-I.T.; visualization, K.K., R.Y., S.Y.; writing (original raft), K.K.; writing (review and editing), all authors.

## Data and Materials availability

All data needed to evaluate the conclusions in the paper are present in the paper and/or the Supplementary Materials.

## Supplementary Materials for

**Fig. S1.**
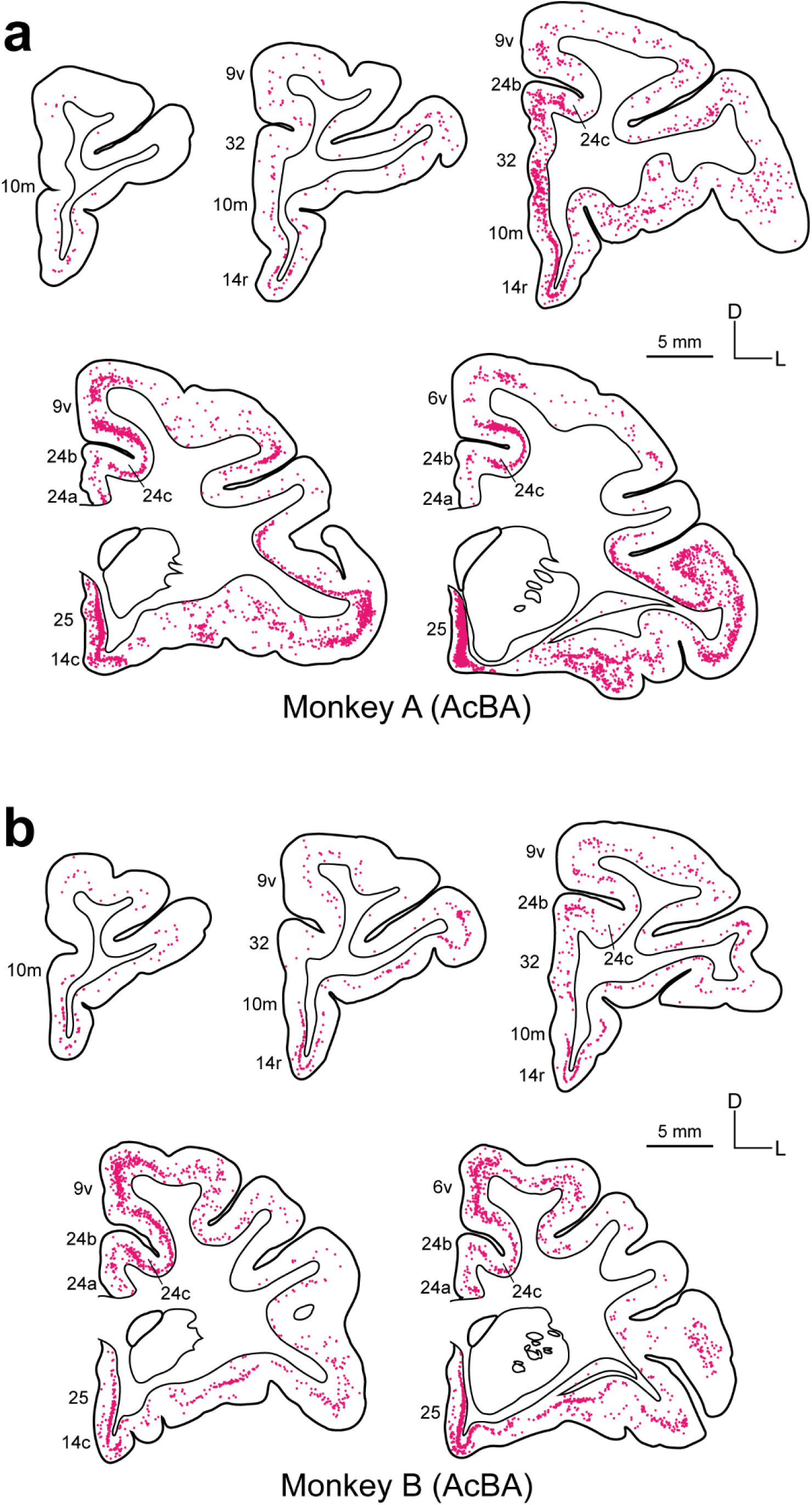
Distribution of retrogradely labeled neurons in the MFC after AAV2retro injections into the AcBA in Monkeys A (a) and B (b) In each panel, shown are five representative coronal sections through the MFC that are arranged anteroposteriorly. Each dot represents one labeled neuron.

**Fig. S2.**
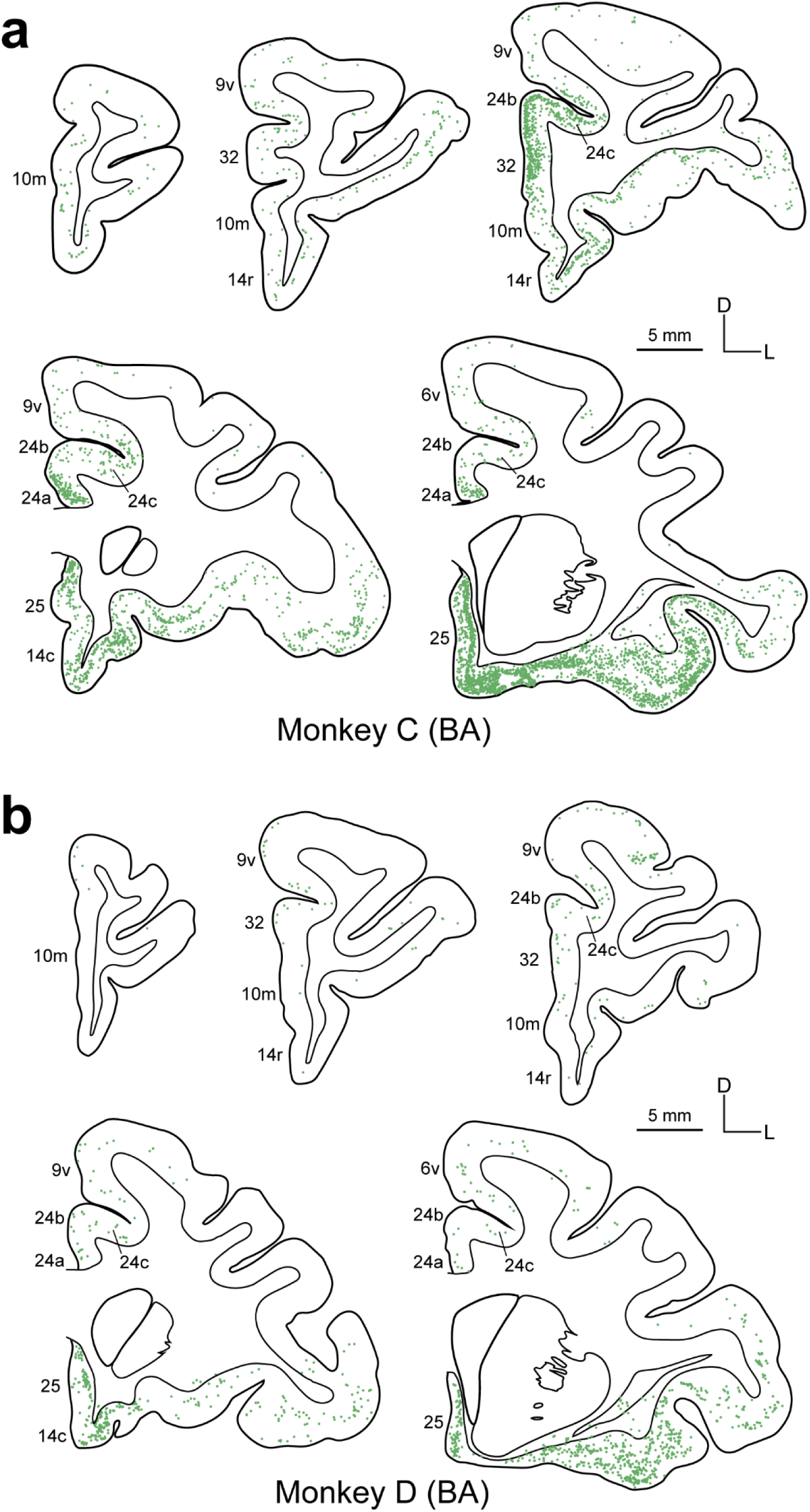
Distribution of retrogradely labeled neurons in the MFC after AAV2retro injections into the BA in Monkeys C (a) and D (b) All conventions are as in Supplementary Figure S1.

